# Systematic search for new HLA alleles in 4195 human 30x WGS samples

**DOI:** 10.1101/2024.05.31.596796

**Authors:** EA Albert, AA Deviatkin, DI Smirnova, M. Woroncow, G.Y Zobkova, A.V Smirnova, PY Volchkov

## Abstract

HLA (Human Leukocyte Antigens) is a highly polymorphic locus in the human genome which also has a high clinical significance. New alleles of HLA genes are constantly being discovered but mostly through the efforts of laboratories which primarily focus on HLA typing and are using field-specific experimental and data processing techniques, like enrichment of HLA region in high-throughput sequencing data. Nevertheless, a vast amount of whole genome sequencing (WGS) data was accumulated over the past years and continues to expand rapidly. Therefore it is an appealing possibility to identify new HLA alleles and refine the information on known alleles from already available WGS data. Currently there are many tools designed for HLA typing, e.g. assigning known alleles, from non HLA enriched WGS data, but none of them specifically tailored towards identification and immediate thorough description of new HLA alleles. Here we are presenting a pipeline HLAchecker, which is specifically designed to identify potentially new HLA alleles based on discrepancies between predicted HLA types, made by any other dedicated tool, and underlying raw 30x WGS data. HLAchecker reports structured in a way which simplifies further validation of potentially new HLA alleles and streamlines submission of alleles to appropriate databases. We validated this tool on 4195 30x WGS samples typed by HLA-HD, discovered 17 potentially new HLA alleles with substitutions in exonic regions and validated five randomly chosen alleles by Sanger sequencing.

## Introduction

The HLA (Human Leukocyte Antigens) genotype is an important feature of the human genome that has various clinical implications, such as susceptibility or resistance to certain diseases and donor - recipient compatibility for a bone marrow transplantation. Therefore, HLA genotyping is widely used in routine clinical practice (1). HLA genotyping is performed by comparing the sample with a set of references. This can be done by polymerase chain reaction (PCR) (2) or high-throughput sequencing (HTS) (3). In the clinical setting, the HTS approach is usually tailored to high coverage sequencing of the HLA locus by using specific HLA enrichment kits. The enrichment of HLA loci for sequencing with specific kits increases the accuracy of the analysis and reduces its price. Therefore, HLA typing in routine clinical practice today is performed either with HLA locus PCR enrichment kits and subsequent sequencing or with a direct PCR test. It should be noted that both methods can be used to determine whether a patient has an allele that is similar to an allele contained in the reference database. If the patient has an HLA haplotype that does not match the allele in the reference set, the analysis will identify the HLA haplotype from the reference set that is most similar to the patient’s allele. Consequently, a new HLA haplotype that is not included in the reference database would not be identified without special efforts and dedicated bioinformatics analysis. Nevertheless, the WGS data collected over the years provide a good opportunity to systematically refine the HLA sequence database, which for historical reasons often contains only partial sequences of alleles, and to identify new HLA alleles.

Over the last fifteen years, a number of tools have been developed to solve the task of HLA genotyping from WGS data(4–7). Such genotyping is a complicated process due to the high similarity between alleles of HLA genes and the sparse nature of the IPD-IMGT/HLA database, in which some alleles are only represented by partial cds sequences. Some of the tools developed could be used in the in-house pipeline to recognize new HLA alleles from WGS data (e.g. kourami and HISATgenotype). But none of these tools are directly tailored for the discovery and characterization of new alleles. Therefore, a lot of manual labor is required to systematically screen large cohorts of samples for potentially new HLA alleles. As far as we know, the only tool specifically suited for this task - NovAT (8) - is only available as a web application, which makes its implementation into the running analysis pipeline impossible and poses problems with sensitive data management.

In the current study, we have developed a freely available tool to characterize HLA alleles based on 30x WGS data - HLAchecker https://gitlab.com/EugeneA/hlachecker. HLAchecker operates over HLA typing results from any of HLA typing tools and WGS aligned bam file. HLA typing results are tested against raw data and a detailed report is generated for each case of discrepancy, which marks a potentially novel allele of the HLA gene. The report is specifically designed to expedite further validation and submission of the discovered allele to appropriate databases. We validated the tool on a cohort of 4195 WGS samples typed with HLA-HD, identified 17 potentially new HLA alleles and validated five new randomly chosen alleles by Sanger sequencing.

## Methods

### Pipeline description and organization

All steps of data analysis are described in the result section of the manuscript. Technically pipeline is written in *wdl* language, which brings all the advantages of specific pipeline oriented frameworks such as logical task based structure, containerisation of individual tasks with docker, resource management and butch submissions. Samtools (9) and Minimap2 (10) are used to extract reads, which fall on the 5MB region of chr6 from WGS bam file and remap these to HLA genes from IPD-IMGT/HLA (11). Custom script written on biopython used to generate reference sequences and supplementary sequences (specified in the main text) for the further usage in the pipeline. Bowtie2 (12) is used to map selected reads to generated references. Bcftools with following parameters “*call -mv -P 1*.*1e-2 --ploidy 1 “* is used to call variants based on the resulting bam files. Custom script written on biopython is used to identify variants in exons and define amino acid substitutions. If variants in exonic regions were detected, blastn (13) is used to search obtained consensus sequences against local copy of Nt db. Mosdepth (14) is used to calculate mean coverage for each allele of each gene. Igv is used to visualize results for candidate cases. All tools are packed in docker containers and uploaded to dockerhub.

### Cohort selection

To evaluate the possibility of discovering new HLA alleles based on standard 30x WGS data, we used samples collected by “Evogen” during routine sequencing of the healthy Russian population. We have selected WGS data for 4195 individuals, with homogeneous genetic background, based on PCA analysis, performed as described previously (15). Homogeneous genetic background was required for correct analysis of HLA haplotypes frequencies.

### Library preparation and sequencing

Library preparation and sequencing was performed as described previously (15). Briefly DNA extraction was performed by spin column using the Qiagen QIAamp DNA Blood Kit (Cat.No. 51106) from whole blood according to the manufacturer’s protocol. DNA amount was measured fluorometrically with Qubit4 (Thermo Fisher Scientific) / Denovix (DeNovix Inc). For the subsequent library preparation only genomic DNA of high quality (OD260/OD280=1.8-2.0, OD260/OD230>2.0) was used. Library preparation was performed with a PCR-free enzyme fragmentation protocol (MGIEasy FS PCR-Free DNA Library Prep Set, Cat.No. 1000013455) using 800-1200 ng gDNA. The distribution of insert size was 400-600 bp. WGS library preparation was performed both manually and automatically. Whole genome sequencing was performed using DNBSEQ-G400 (MGI Tech Co., Ltd.) with FCL PE150 (cat. no. 1000012555), FCL PE200 (cat. no. 1000013858), and DNBSEQ-T7, according to the manufacturer’s protocol. Average sequencing depth was 30x.

### Sanger validation

Randomly chosen 5 samples with detected new allele candidates were validated by Sanger sequencing. Two pairs of allele specific PCR primers were chosen for each sample with ncbi primer-blast software (Supplementary table 1). Amplified PCR fragments were extracted from agarose gel Supplementary figure 1 and sent for Sanger sequencing to Evrogene.

### HLA typing and haplotype frequencies

HLA typing from WGS was performed using HLA-HD (16) according to authors recommendations. HLA haplotypes frequencies were calculated using Hapl-o-Mat according to authors recommendations.

### Ethics

The study was approved by the local ethical committee of the Independent Multidisciplinary Committee on Ethical Review for Clinical Trials (Moscow, Russia) and was performed in accordance with the approved guidelines and the Declaration of Helsinki.

## Results

### Structure of the pipeline for identification of new HLA alleles from WGS data

The overall structure of the pipeline to identify potential new alleles of HLA genes that have variations in exonic sequences is shown in Figure 1A. This pipeline uses the results of whole human genome sequencing available in BAM or CRAM (Binary Alignment Map) format and HLA typing results from any of the dedicated tools. Only reads mapping to the 5Mb locus of the sixth chromosome as well as to the alternative contigs of the sixth chromosome, which contains the HLA locus, were used for further processing (step 1 at Figure 1A). Filtered reads were mapped to all possible reference genomic sequences from the IPD-IMGT/HLA database (step 2 at Figure 1A). Only reads mapped to selected HLA genes (DQA1, DQB1, DRB1, A, B, C) were used for further analysis (step 3 at Figure 1A). These reads were used for mapping to the genomic sequence of the alleles for each given gene predicted by the external HLA typing tool (step 4 at Figure 1A). In the current study, HLA-HD was used for typing HLA for both alleles in each sample.

**Figure 1.**
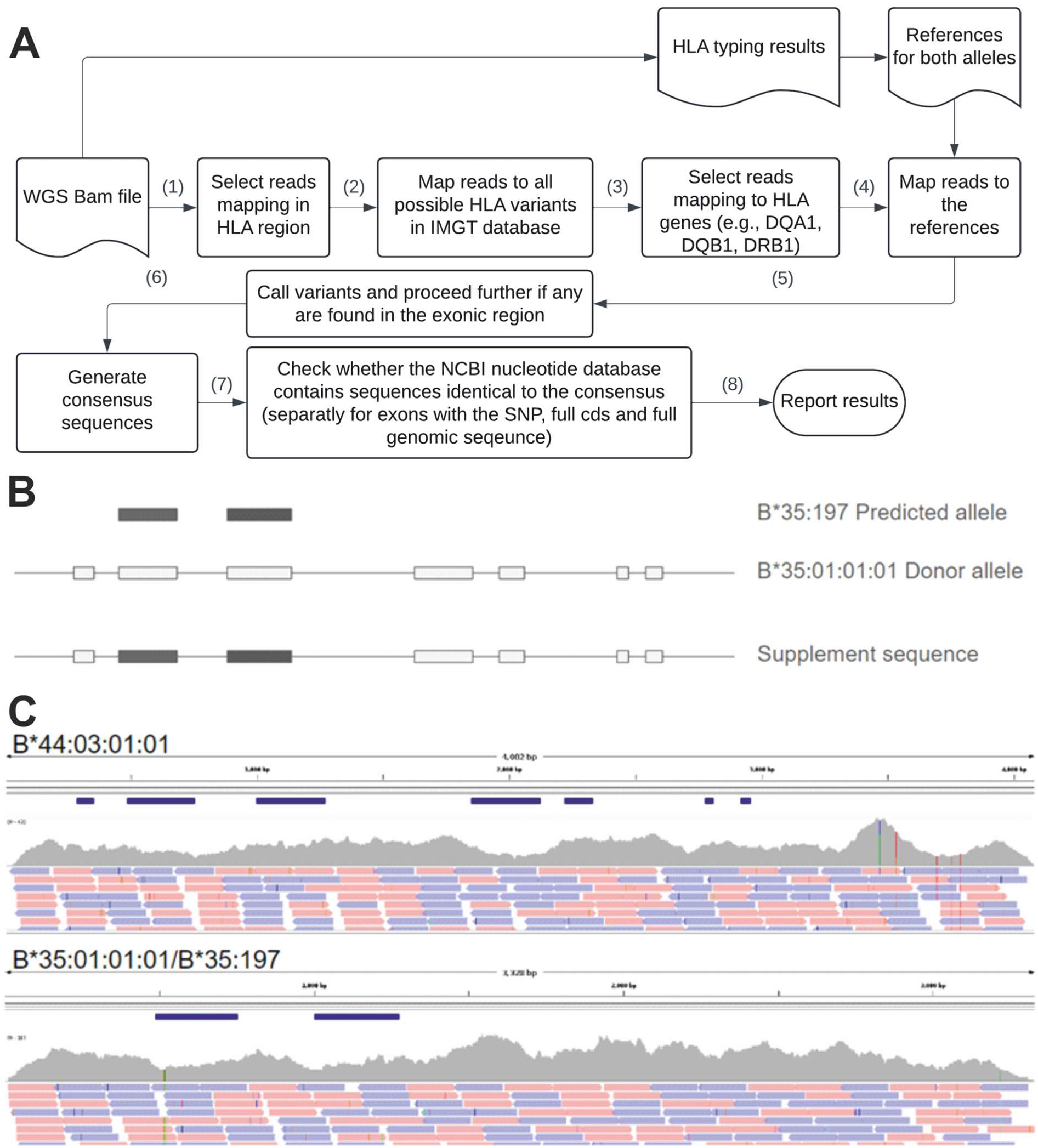
Schematic pipeline representation. A. Overall pipeline schema. B Example of supplement sequence generation for allele which lack full gene sequence. C. Example reads mapped to gene B for a sample with defined haplotypes B*44:03:01:01/B*35:197, where absent genomic sequence for B*35:197 was supplemented with B*35:01:01:01. Coverage, reads and exonic regions track are shown. Notice two SNPs in B*35:197 exon.

It should be noted that for some HLA alleles the IMGT database contains only sequences of individual exons. In these cases, HLAchecker creates a supplementary sequence with the missing part of the HLA allele. The artificial sequence is created by supplementing the missing part of the allele of interest with genomic sequences of the closest allele based on the HLA nomenclature (Figure 2B).

**Figure 2.**
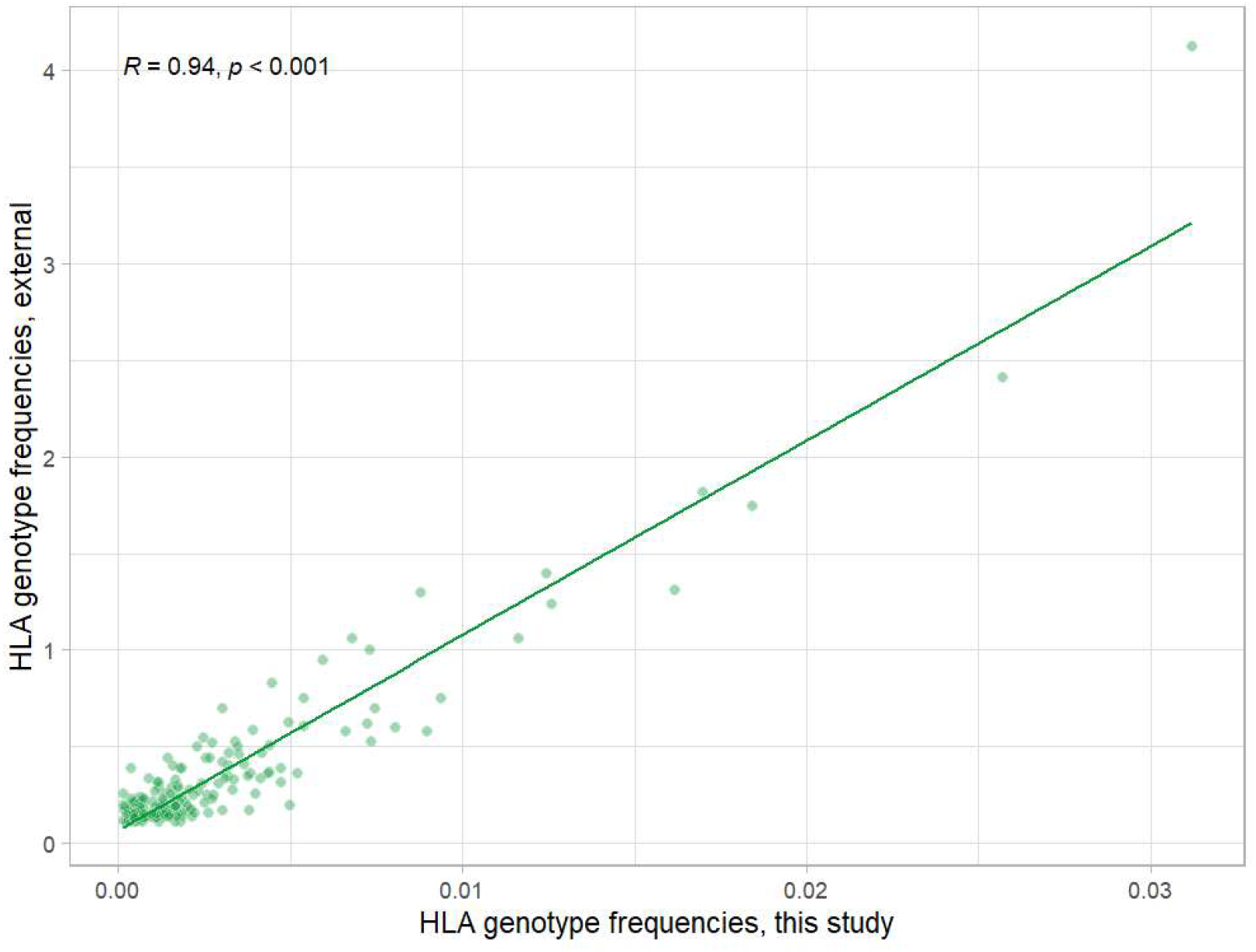
Correlation of frequencies for A-B-DRB1 haplotypes between data from www.allelefrequencies.net for central russian population with frequencies from current studies (17).

The mapping (step 4 at Figure 1A) was performed separately for each gene and a set of selected reads. Both alleles of a selected gene were used simultaneously as reference. This approach allows automatic phasing of alleles (Figure 1C). The obtained alignment files were filtered for reads with more than 2 mismatches from the reference, to filter highly diverged reads, presumably originated from pseudogenes which are known to exist at least for some HLA genes. The generated bam files were used for SNP/INDELS calling (step 5 at Figure 1A). If any variants are called in the exonic region of HLA allele then further analysis is conducted. First of all genomic consensus sequences were generated for the allele with the discovered variant. At that stage both exonic and intronic variants were used (step 6 at Figure 1A). It should be noted that the IMGT database is not updated in real time. Therefore, it is possible to identify sequences that are not included in the latest version of IMGT but are already described in another database. Therefore an additional comparison of the obtained consensus with the less stringent NCBI Nucleotide Database was performed (step 7 at Figure 1A).

NCBI Nucleotide Database contains partial alleles along with full alleles, therefore blastn was performed separetly for exon with the SNP, full cds consensus sequence and full genome consensus sequence. At the last step HLAchecker generated reports with information specifically tailored for submission of discovered alleles to GenBank and IMGT. In addition, screenshots of the mapping of reads to references are generated and visualized with the Integrative Genomics Viewer for each allele with exonic variation to allow easy manual review (step 8 at Figure 2A).

### Developed pipeline successfully identifies discordance between sequences of predicted alleles and WGS data

The purpose of the HLAchecker is to identify and automatically characterize discordance between raw WGS data and the prediction of the HLA typing tool for a given sample. To simulate such discordance and estimate pipeline efficiency we introduced random exonic SNPs in reference sequences of HLA allele prior to mapping selected reads with bowtie2 (Figure 1A, step 4). We selected 100 random samples from the cohort where the initial run of the pipeline did not identify any HLA associated mismatches. We repeated the analysis and introduced one random exonic SNP in the reference sequence of each HLA allele for each sample for 6 main HLA genes: A,B,C, DQA1, DQB1, DQR1. Pipeline successfully identified 1105 out of 1135 artificially introduced SNPs demonstrating that 30x WGS data on average harbor enough reads to correctly identify samples with new HLA alleles. Therefore the sensitivity of HLAcheker was estimated to be 97.4% (Table 1). It should be noted that such simulation design does not allow to calculate the number of false positives and true negatives.

**Table 1.** Contingency table demonstrating HLAchecker sensitivity.

|  | **Predicted Positive** | **Predicted negative** |
| --- | --- | --- |
| Actual positive | True positive = 1135 | False negative = 30 |
| Actual negative | False positive = N/A | True negative = N/A |

We manually examined the SNPs that were missed by HLAchecker. They were all located in the regions that are identical between two alleles, resulting in a mapping of the reads to the paired unmodified reference sequence (Supplementary Figure 2A, 2B).

### HLA allele frequencies in the cohort used for HLAchacker validation were consistent with available populational data

To validate HLAchecker on the real 30x WGS data we selected a cohort of people from a dataset of LCC Evogene, which was obtained through routine 30x WGS of healthy individuals. A population analysis and filtering were performed to prevent a possible enrichment of the cohort with the new alleles and the associated bias. The cohort was restricted to individuals of Central Russian ancestry, which is well characterized in terms of the frequency of HLA alleles. In total, the genomic information of 4195 individuals was used (Supplementary Figure 3). HLA typing of the cohort was performed using the HLA-HD tool and subsequent analysis was restricted to the major HLA genes: A,B,C, DQA1, DQB1, DQR1. To validate the obtained HLA-HD results with the orthogonal method, we compared the frequencies of haplotypes in the selected cohort with the data from the www.allelefrequencies.net database for the Moscow population (Fig. 2, supplementary table 2). Haplotype frequencies were calculated with the Hapl-o-Mat software, according to guidelines. Overall, the results of HLA typing with HLA-HD showed a good correlation with the independent dataset created using a different method. Therefore, the results obtained from our cohort can be used for further validation of HLAchecker.

### Analysis of the 4195 30x WGS samples revealed 17 of potentially new HLA alleles

To test the capability of HLAchecker to identify new alleles in 30x WGS data we run it on the aforementioned cohort. A statistical summary of the results can be found in Table 2. The frequency of previously unknown exon sequences varied for the six major HLA genes, ranging from 0.00024 for B and C to 0.00071 for DQA1. The higher number of missing protein coding and genomic sequences in the database clearly shows that for some HLA alleles no complete genomic or exonic sequences are available.

**Table 2.** Summary of analysis of 4195 samples for discrepancies between sequences of predicted HLA alleles and WGS data. Selected samples with exonic variants coverage above 5.

|  | The number of samples with potential new allele | The number of unique potential new alleles | The number of unique potential new alleles with exonic sequence with variant absent in the NCBI Nucleotide database | The number of unique potential new alleles with protein coding sequences absent in the NCBI Nucleotide database | The number of unique potential new alleles with full genomic sequence absent in the NCBI Nucleotide database |
| --- | --- | --- | --- | --- | --- |
| A | 143 | 103 | 2 | 7 | 17 |
| B | 96 | 81 | 2 | 17 | 33 |
| C | 95 | 90 | 3 | 20 | 32 |
| DQB1 | 12 | 6 | 2 | 2 | 3 |
| DQA1 | 29 | 17 | 6 | 11 | 12 |
| DRB1 | 40 | 22 | 2 | 11 | 19 |
| TOTAL | 415 | 319 | 17 | 68 | 116 |

Sanger sequencing is a golden standard for validation of any alteration in DNA, therefore a set of variants identified in cohort by the pipeline were validated by Sanger sequencing. We randomly selected five samples with the exonic sequences absent from NCBI Nucleotide Database for validation (Table 3). For each sample two pairs of allele specific primers were selected which selectively amplify either the exonic region from the allele with the variant or the corresponding region from the non-altered allele (Supplementary figure 1, Supplementary table 1). Sanger sequencing of PCR products was performed in both directions from the same primers according to the submission guidelines of the IPD-IMGT/HLA database. All 5 new allelic variants were validated through Sanger sequencing and one of them was submitted to the IMGT database. Sanger sequencing results are summarized in table 3.

**Table 3.** Summary of pipeline prediction validations with sanger sequencing.

| Sample | Allele | CDS position | Substitution in the nucleotide | Substitution in the protein | Exon |
| --- | --- | --- | --- | --- | --- |
| 1 | HLA00613 | 169 | C -> A<br>(nonsynonymous) | Q -> K | 2 |
| 2 | HLA00434 | 828 | A -> C<br>(synonymous) | - | 4 |
| 3 | HLA00723 | 567 | G -> A<br>(nonsynonymous) | M -> I | 3 |
| 4 | HLA00756 | 429 | C -> T<br>(synonymous) | - | 3 |
| 5 | HLA00601 | 672 | G -> A<br>(synonymous) | - | 3 |

## Discussion

In this paper we demonstrated the possibility of identification of new HLA alleles from standard 30x WGS human sequencing samples and presented the pipeline specifically tailored for that purpose. Such approach can help to expand HLA allele repertoire by reanalysis of HLA region in wide cohorts of existing whole genome sequencing (e.g. Gnomad (18) and UK biobank (19)). Such an approach can help to fill the gaps in the IPD-IMGT/HLA database supplementing missing full genomic sequences which pose a serious problem in the HLA analysis. It has to be noted, that even though reanalysis of accumulated short reads 30x WGS could provide a good way of high throughput screening for new alleles it does not have 100% accuracy in all cases and independent validation (e.g. Sanger sequencing) will be required for identified candidates.

Several limitations are associated with the proposed approach. First of all it relies on HLA typing performed by some external tools. False positive calls may arise from inaccurate typing or the version mismatch between the IPD-IMGT/HLA database used for typing and used in the pipeline. Some individuals harbor pseudogenes in their genome, and reads from these regions might interfere with the HLAchecker processing, as well as with upstream HLA typing tools. Due to the sparse nature of the IPD-IMGT/HLA and presence of substantial amounts of alleles without full genomic sequences, we choose not to report variation in the intronic regions which limits application of our approach to the third field of HLA nomenclature.

We reanalysed HLA in a cohort of 4195 individuals, which originated from central Russia to estimate frequency of new alleles in that relatively well characterized population. In total we identified 319 alleles of 6 genes which had mismatches with predicted sequences from IPD-IMGT/HLA and the coverage distinguishing SNP above 5. We separately performed additional searches of the identified exons with variants 17 exonic, 68 complete protein coding and 116 complete gene sequences were absent from the NCBI Nucleotide database. Therefore even in a relatively well characterized population which is genetically similar to Europeans reanalysis of a 4195 individuals cohort yields a substantial amount of valuable information.

One of the main problems with HLA analysis tools is the need to constantly update the database. Minor changes in the database format cause scripts that generate input for the tools to become obsolete. For example, launching a popular tool for HLA typing - Kourami (4), with the newest IPD-IMGT/HLA release requires manual correction of the files. Pipeline presented here takes raw fasta files for complete protein coding and gene sequences from IPD-IMGT/HLA which provide a most convenient way of keeping data up to date for the analysis. Moreover all steps of the pipeline are containerised with docker, making it robust and easy to launch.

Over the years certain guidelines were developed for registry of the new HLA alleles. Pipeline, presented in the current publication specifically tailored to providing all necessary information on detected event e.g. new allele consensus sequence, position of variants in the genomic sequence, amino acid changes, number of exon affected by the variants and results of blast analysis of consensus sequence against Nt database. In total that information substantially simplifies following steps of the allele characterisation - Sanger sequencing validation, submission to gene bank and submission to IPD-IMGT/HLA.

## Data availability

Code of the described pipeline available at the github https://gitlab.com/EugeneA/hlachecker.

## Supplementary materials

**Supplementary figure 1.**
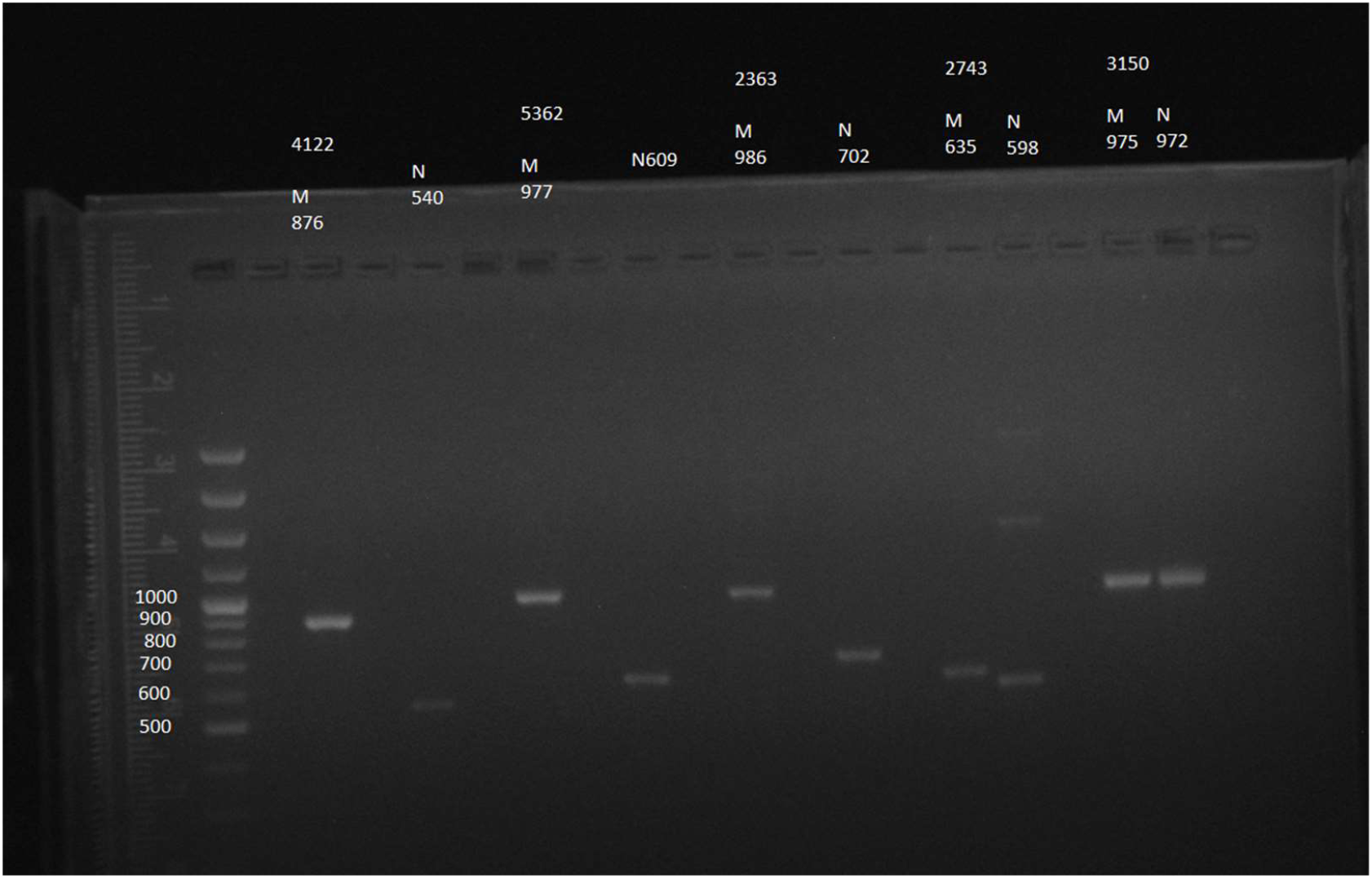
Allele specific PCR of five samples for fragments generation and subsequent Sanger sequencing.

**Supplementary table 1.**
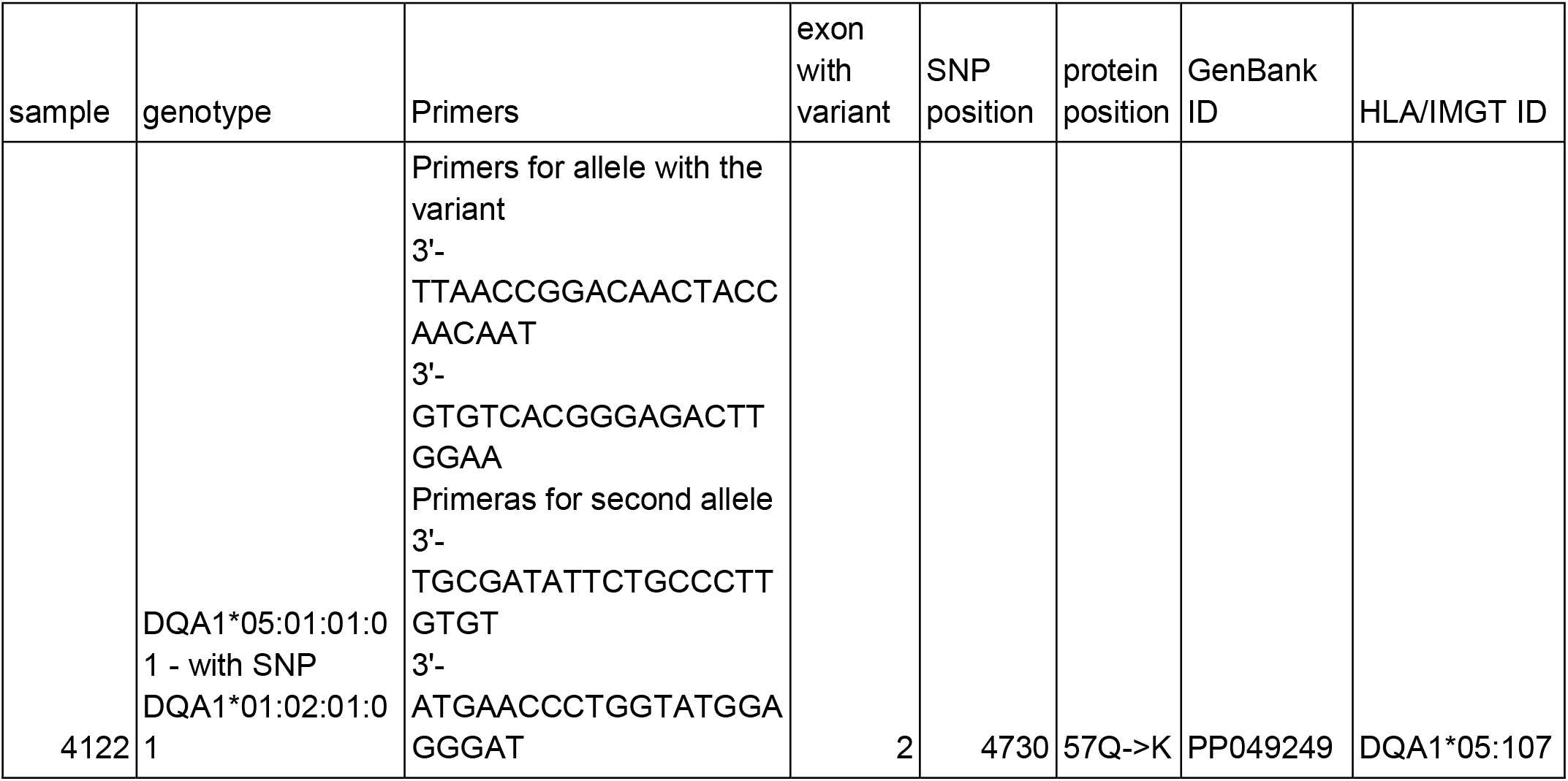
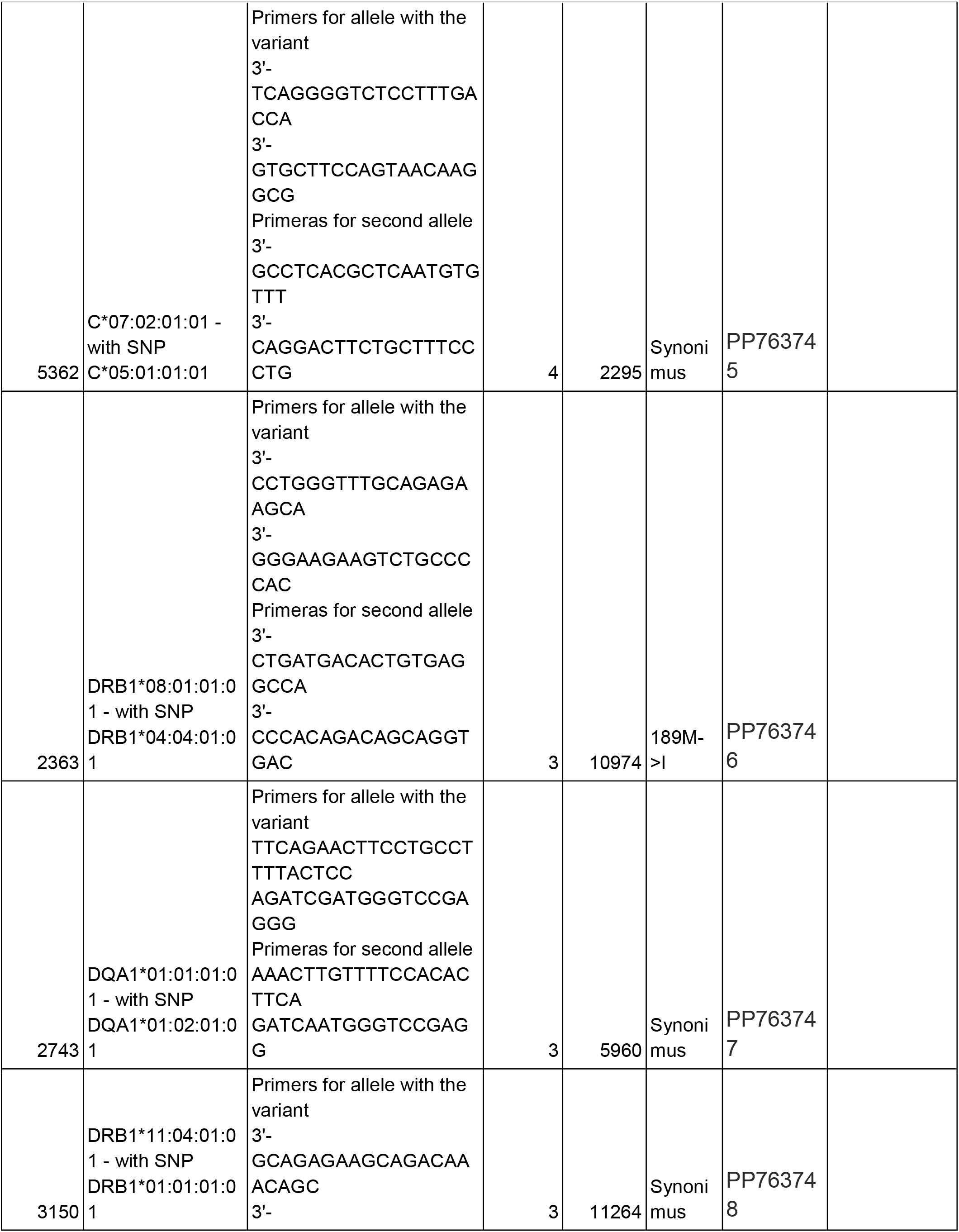
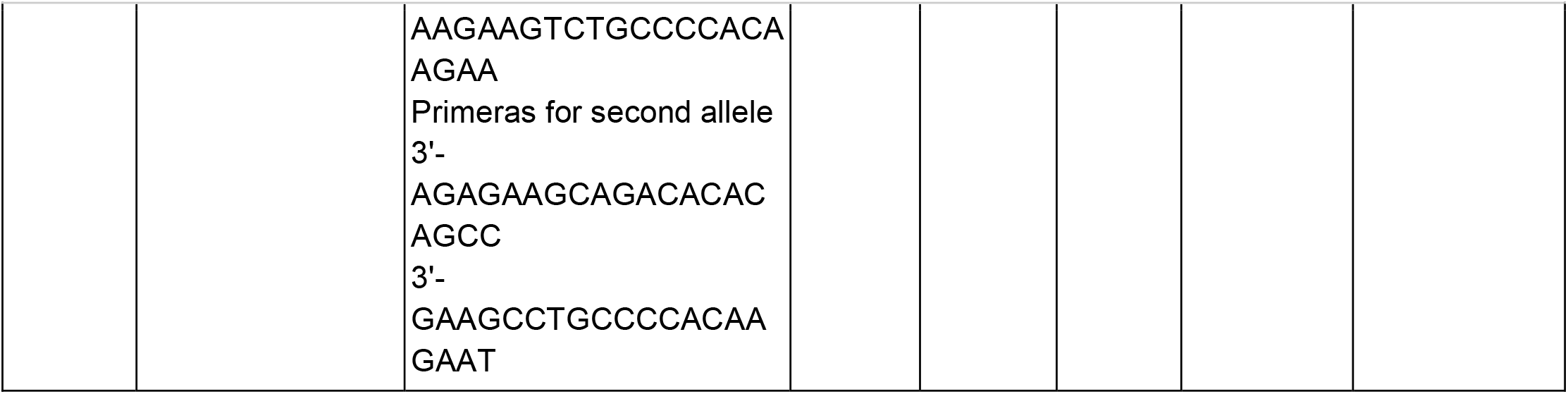
Sanger validation of 5 candidate variants.

**Supplementary figure 2.**
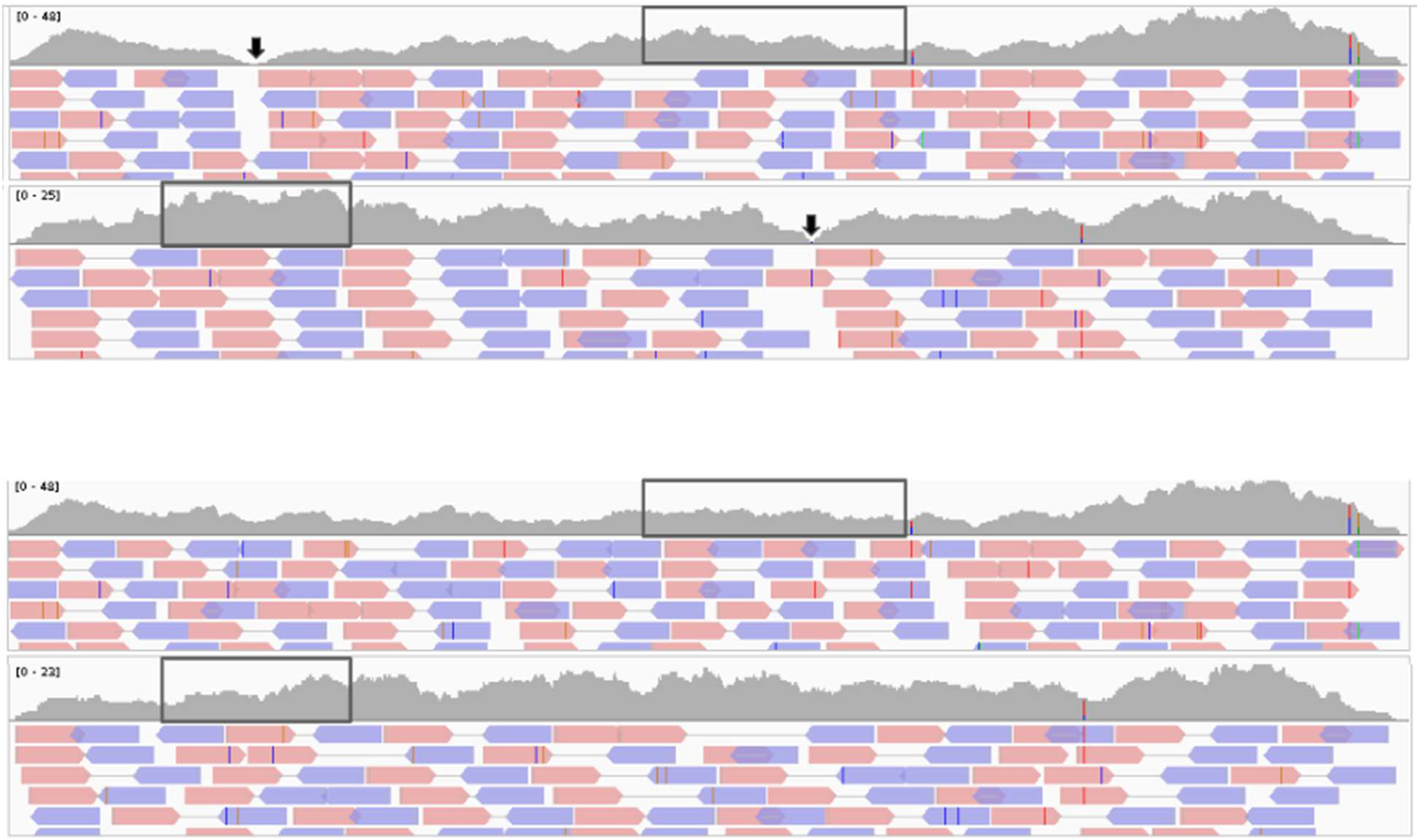
Example of false negative call for HLAchecker during in-silico validation. A. Mapping to both alleles of C gene is present. Random SNPs positions are marked with an arrow.. B. Same sample as on A but no random SNPs were introduced. Note two gaps in the coverage around SNPs and changes in boxed coverage of corresponding alleles between A and B.

**Supplementary figure 3.**
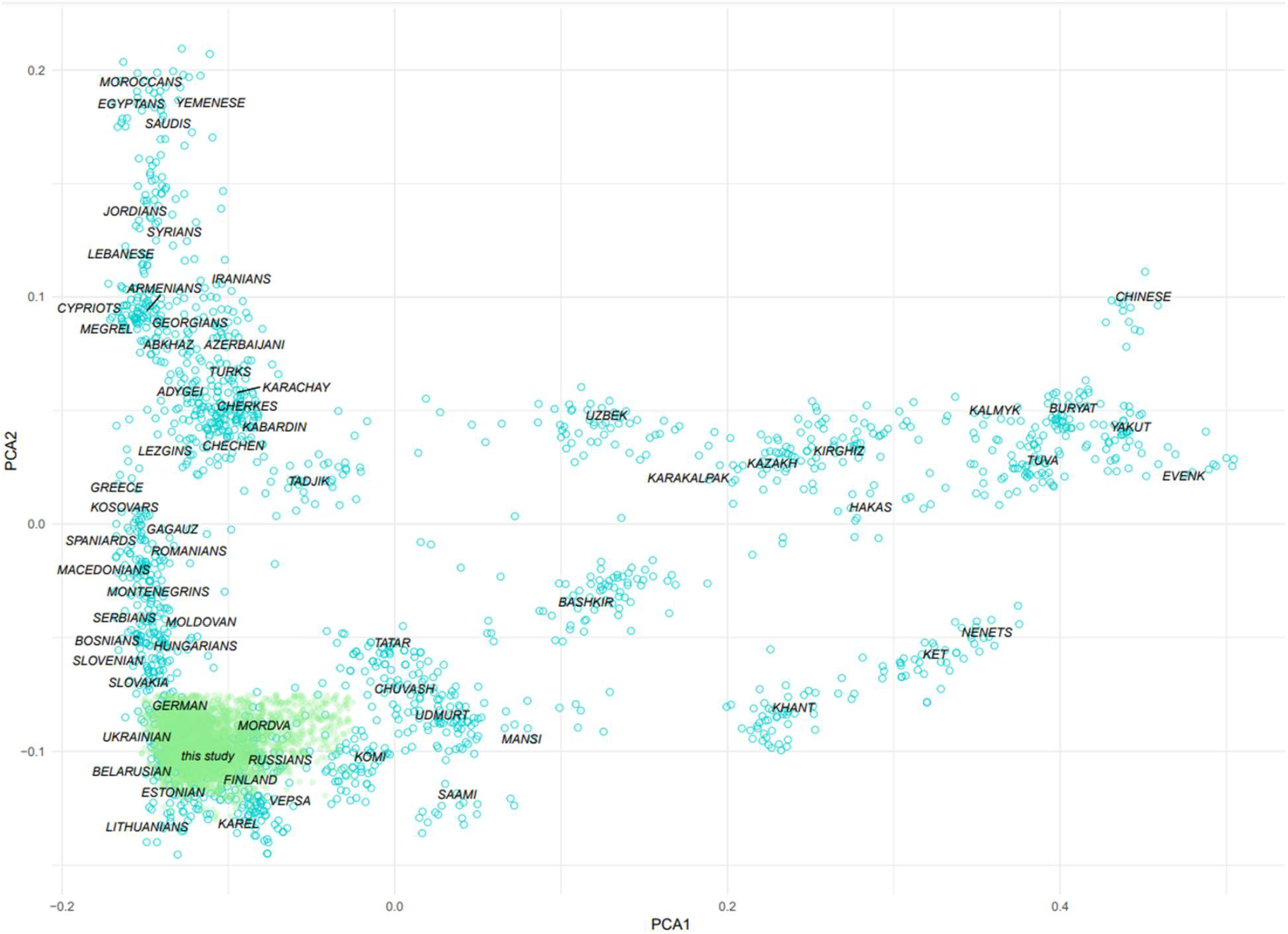
Ethnic analysis of the cohort used in the study. Blue circles represent the reference population, green circles represent cohort from the current study. PCA is built on the 400000 SNPs.

